# Highly specialized carbohydrate metabolism capability in *Bifidobacterium* strain associated with intestinal barrier maturation in early preterm infants

**DOI:** 10.1101/2022.05.06.490995

**Authors:** Bing Ma, Sripriya Sundararajan, Gita Nadimpalli, Michael France, Elias McComb, Lindsay Rutt, Jose M Lemme-Dumit, Elise Janofsky, Lisa S. Roskes, Pawel Gajer, Li Fu, Hongqiu Yang, Mike Humphrys, Luke J Tallon, Lisa Sadzewicz, Marcela F Pasetti, Jacques Ravel, Rose M Viscardi

**Author notes:** Address correspondence to Bing Ma.

## Abstract

“Leaky gut”, or high intestinal barrier permeability, is common in preterm newborns. The role of microbiota in this process remains largely uncharacterized. We employed both short- and long-read sequencing of the 16S rRNA gene and metagenomes to characterize the intestinal microbiome of a longitudinal cohort of 113 preterm infants born between 24^0/7^-32^6/7^ weeks of gestation. Enabled by enhanced taxonomic resolution, we found significantly increased abundance of *Bifidobacterium breve* and a diet rich in mother’s breastmilk to be associated with intestinal barrier maturation during the first week of life. We combined these factors using genome-resolved metagenomics and identified a highly specialized genetic capability of the *Bifidobacterium* strains to assimilate human milk oligosaccharides and host-derived glycoproteins. Our study proposed mechanistic roles of breastmilk feeding and intestinal microbial colonization in postnatal intestinal barrier maturation; these observations are critical towards advancing therapeutics to prevent and treat hyperpermeable gut-associated conditions, including necrotizing enterocolitis.

**IMPORTANCE:** Despite improvements in neonatal intensive care, necrotizing enterocolitis (NEC) remains a leading cause of morbidity and mortality. “Leaky gut”, or intestinal barrier immaturity with elevated intestinal permeability, is the proximate cause of susceptibility to NEC. Early detection and intervention to prevent leaky gut in “at-risk” preterm neonates is critical to lower the risk for potentially life-threatening complications like NEC. However, the complex interactions between the developing gut microbial community, nutrition, and intestinal barrier function, remain largely uncharacterized. In this study, we revealed the critical role of sufficient breastmilk feeding volume and specialized carbohydrate metabolism capability of *Bifidobacterium* in coordinated postnatal improvement of intestinal barrier. Determining the clinical and microbial biomarkers that drive the intestinal developmental disparity will inform early detection and novel therapeutic strategies to promote appropriate intestinal barrier maturation, prevent NEC and other adverse health conditions in preterm infants.

---

Early preterm neonates are particularly vulnerable to life-threatening events and routinely require intensive care and medical intervention to survive (1). The physiological immaturity of their gastrointestinal (GI) tract is commonly associated with deficiencies in barrier functions that result in a clinical syndrome known as “leaky gut” (2–5). Under leaky gut condition, the bacteria and bacterial products normally confined to the intestinal lumen are able to translocate into the peripheral circulation through the hyperpermeable epithelial barrier, which could lead to widespread invasion of the intestinal epithelium and gut lamina propria, mucosal inflammation, epithelial cell damage, intestinal necrosis, systemic infection, and ultimately multi-organ failure and death (4, 6, 7). Necrotizing enterocolitis (NEC) is a prominent bacterial translocation-associated GI condition that affects 7-10% of preterm neonates or 1-5% of all neonatal NICU admissions with a devastating mortality rate as high as 50% (8–12). Early detection of an aberrant leaky gut and early intervention to limit intestinal injury are of paramount importance to reduce the incidence of subsequent complications including NEC (12, 13).

A functional intestinal barrier combines a physical barrier that encompasses chemical, immunological and microbiological components (14). We and others have found that the first week of life (day 8±2 post-birth) is a critical window during which the most rapid postnatal intestinal maturation occurs (15–17). More importantly, these earlier studies demonstrated that the intestinal barrier function, which develops mostly *in utero* in term infants, can be improved postnatally. They also showed that the intestinal barrier maturation does not occur at the same rate, with ∼40% of preterm neonates (<33 weeks gestation) failing to develop a functional intestinal barrier within the first two weeks of life (15, 16). Determining the factors that drive this developmental disparity will inform early detection and novel therapeutic strategies to promote intestinal barrier maturation.

Efforts to characterize the microbiological factors that are associated with intestinal barrier maturation have thus far yielded unsatisfactory results (18). There are no microbial biomarkers predictive of intestinal development. A major limitation is the use of partial 16S rRNA gene sequences to evaluate the taxonomic composition of gut microbiota. The short sequences lack the phylogenetic signal necessary to describe taxonomic composition at species or even genus level. Many of the PCR primers used to amplify variable regions of the 16S rRNA gene fail to amplify members of the genus *Bifidobacterium* (19–21). *Bifidobacterium* species are known to be frequent colonizers of infant guts (22), and are considered to play beneficial roles in intestinal development and influence maturation of the neonatal gut, potentially through stimulating colonic epithelial proliferation, modulation of host defense responses and protection against bacterial infections (23, 24). To investigate *Bifidobacterium* and other bacterial groups predictive of early intestinal development and maturation are of pivotal importance.

In this study, we sought to characterize the role of early assembly of infant gut microbiota and its metabolism in postnatal intestinal barrier maturation. We build upon the results of past studies (15, 16) using an expanded cohort (N=113) of early preterm neonates (24^0/7^-32^6/7^ weeks of gestation) from whom stool samples were collected daily up to 21 days post birth. High resolution approaches were applied to characterize the composition of the developing gut microbiota with substantially enhanced taxonomic resolution including *Bifidobacterium* species, which we identified as the microbial biomarker associated with postnatal intestinal barrier maturation within the first week of life. Whole community metagenomes using both short- and long-read sequences provided a detailed characterization of the genetic content of these *Bifidobacterium* species, which were shown to have distinct genetic features affording complete carbohydrate foraging capabilities, including human milk oligosaccharides (HMOs) and host-derived glycoprotein. The presence of specific strains of *Bifidobacterium* may inform the early detection of aberrant intestinal permeability. Supplementation of these bifidobacterial strains could be leveraged in novel intervention strategies for the prevention of leaky gut and its devastating sequelae in preterm newborns.

## RESULTS

### Clinical cohort

We examined a prospective cohort of 113 preterm infants 24^0/7^-32^6/7^ weeks of gestation including 37 subjects described in a previous analysis (**Table S1**). Fecal samples were collected daily until postnatal day 21 or discharge from the Neonatal Intensive Care Unit (NICU, **Fig. 1**). Mean gestational age (GA) of infants at birth was 29.9±2.3 weeks. A total of 28 infants (24.8%) were <28 weeks GA, and 85 (75.2%) were 28^0/7^-32^6/7^ weeks GA. The mean birth weight was 1,381g (±415g); 66 (58.4%) newborns were classified as very low birth weight (VLBW, <1,500g birth weight) and 26 (23.0%) were classified as extremely low birth weight (ELBW, <1,000g).

**FIG 1.**
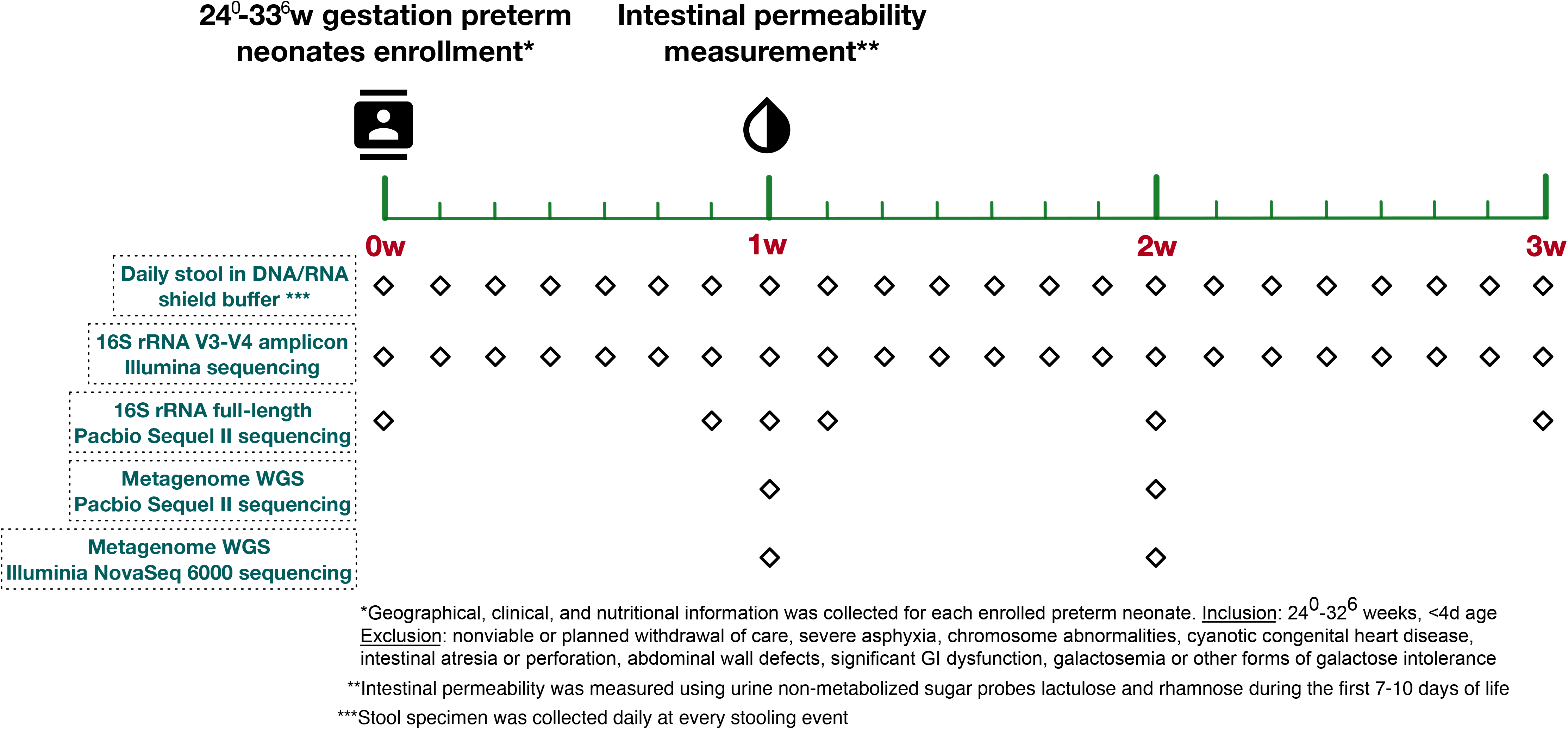
Study design. *Demographic, clinical, and nutritional information was collected for each enrolled preterm neonate. Inclusion: 24^0^-32^6^ weeks, <4d age. Exclusion criteria include nonviable or planned withdrawal of care, severe asphyxia, chromosome abnormalities, cyanotic congenital heart disease, intestinal atresia or perforation, abdominal wall defects, significant GI dysfunction, galactosemia or other forms of galactose intolerance. **Intestinal permeability was measured using urine non- metabolized sugar probes lactulose and rhamnose day 7-10 post-birth. ***Stool specimens were collected daily at every stooling event, stored in storage buffer and archived in -80°C.

Intestinal permeability (IP) was determined 7-10 days post-birth when rapid intestinal barrier maturation normally takes place (15, 16). IP was calculated as the ratio of two enterally administered sugar probes Lactulose (La) and Rhamnose (Rh), markers of intestinal paracellular and transcellular pathways, respectively (25, 26). IP was ranging between 0.001 and 0.394 with an average of 0.07±0.007 (mean±s.e.) and is not significantly different among postnatal day 7-10 (**Supplemental Fig. 1A**). High IP was defined by a La/Rh ratio >0.05, as validated and applied previously (16). Of the 113 subjects, 48 (42.5%) were found to have high IP. Infants <28 weeks GA were more likely to have elevated IP (N=18) than infants 28^0^-32^6^ weeks GA [(64.3% vs. 35.3%), P<0.01].

### Postmenstrual age and mother’s own breastmilk (MOM) feeding are associated with intestinal permeability in early preterm neonates

Among the collected demographic and maternal variables for each infant, four host factors were observed to be inversely related to IP, including: GA, postmenstrual age (PMA) corresponding to chronological and GA, birthweight, and 1-minute Apgar score (**Table 1**). These variables are also highly correlated to one another with high covariates multicollinearity (variance inflation factor > 10) (**Fig. S1**). PMA was the most significant factor associated with IP among the four (P = 0.01, q value = 0.015) based on Hilbert-Schmidt Independence Criterion (HSIC) (**Table S2**). Other host factors such as sex and race were not significantly associated with IP. Maternal factors including preterm premature rupture of membranes (PPROM), maternal antibiotics, antenatal corticosteroids, preeclampsia and delivery mode, were not associated with IP. These data indicate that younger infants have significantly higher incidences of high IP, likely attributed to their more immature intestinal development.

**TABLE 1.** Study cohort demographics and clinical variables stratified by intestinal permeability (IP) category

| Variables | Total cohort<br>(N=113)<br>(N (%) / mean ± SD) | High IP<br>(n=48)<br>(n (%) / mean ± SD) | Low IP<br>(n=65)<br>(n (%) / mean ± SD) | p-value* | <sup>a</sup>Variable measured during the time-period starting from enrollment day (within 1 to 4 days after birth depending on clinical stability) until the day when IP was measured (day 8±2 post-birth). |
| --- | --- | --- | --- | --- | --- |
| Sex |  |  |  | 0.28 |  |
| Male | 61 (54.0) | 24 (50.0) | 37 (57.0) |  |  |
| Female | 52 (46.0) | 24 (50.0) | 28 (43.1) |  |  |
| Race |  |  |  |  |  |
| White | 42 (37.2) | 18 (37.5) | 24 (37.0) | 1.00 |  |
| African American | 63 (55.8) | 30 (62.5) | 33 (50.8) | 0.25 |  |
| Other | 8 (7.1) | 0 | 8 (12.3) | 0.02 |  |
| Birth weight (gram) | 1377.8 ± 415.2 | 1237.3 ± 378.1 | 1496.5 ± 403.0 | <0.01 | |
| VLBW (<1,500 g) | 66 (58.4) | 32 (66.7) | 34 (52.3) | 0.18 |  |
| Gestational age (wks) | 29.8 ± 2.3 | 29.0 ± 2.3 | 30.5 ± 2.1 | <0.01 | |
| Early GA (≤28 wks) | 28 (24.8) | 18 (37.5) | 10 (15.4) | <0.01 | |
| Postmenstrual age (wks) | 31.1 ± 2.3 | 30.3 ± 2.3 | 31.7 ± 2.1 | <0.01 | |
| Early PMA (<31 wks) | 41 (36.3) | 23 (47.9) | 18 (27.7) | 0.03 |  |
| Caesarean delivery | 77 (68.1) | 37 (77.1) | 40 (61.5) | 0.10 |  |
| PPROM | 36 (31.9) | 15 (31.3) | 21 (32.3) | 1.00 |  |
| Preeclampsia | 25 (22.1) | 11 (23.0) | 14 (21.5) | 1.00 |  |
| Antenatal corticosteroids | 106 (94.0) | 46 (96.0) | 60 (92.3) | 0.70 |  |
| Maternal antibiotics | 69 (61.1) | 30 (62.5) | 39 (60.0) | 0.85 |  |
| APGAR Score at 1 min | 5.8 ± 2.5 | 5.3 ± 2.8 | 6.2 ± 2.1 | 0.04 | |
| APGAR Score at 5 min | 7.7 ± 1.6 | 7.5 ± 1.9 | 7.9 ± 1.6 | 0.12 | |
| Antibiotic types |  |  |  |  |  |
| Ampicillin received | 64 (56.7) | 30 (62.5) | 34 (52.3) | 0.33 |  |
| Gentamycin received | 56 (49.6) | 25 (52.1) | 31 (47.7) | 0.70 |  |
| Vancomycin received | 8 (7.1) | 6 (12.5) | 2 (3.1) | 0.07 |  |
| Cefotaxime received | 9 (8.0) | 6 (12.5) | 3 (4.6) | 0.16 |  |
| Received at least one antibiotic vs. no antibiotics<sup>a</sup> | 68 (60.2) | 33 (68.8) | 35 (53.9) | 0.12 |  |
| Antibiotics days received<sup>a</sup> |  |  |  |  |  |
| ≤ 3 days | 83 (73.5) | 30 (62.5) | 53 (81.5) | 0.03 | |
| > 3 days | 30 (26.6) | 18 (37.5) | 12 (18.5) |  |  |
| Days received MBM<sup>a</sup> |  |  |  |  |  |
| < 4 days | 26 (23.0) | 20 (41.7) | 6 (9.2) | < 0.01 |  |
| ≥ 4 days | 87 (77.0) | 28 (58.3) | 59 (90.8) | | |
| Feeding duration (number days)<sup>a</sup> |  |  |  |  |  |
| Mother's own breast milk | 4.8 ± 2.3 | 4 ± 2.7 | 5.5 ± 1.5 | <0.01 | |
| Formula | 1.3 ± 2.3 | 2 ± 2.7 | 0.8 ± 1.6 | 0.02 | |
| Feeding intake volume received<sup>a</sup> |  |  |  |  |  |
| Mother's own breast milk | 200.8 ± 178.8 | 123.4 ± 154.2 | 263.0 ± 175.6 | < 0.01 | |
| Formula | 61.7 ± 146.7 | 99.8 ± 194.7 | 32.8 ± 91.2 | 0.03 | |

However, host factors could only partially explain IP. Mother’s own breastmilk (MOM) longer feeding and higher intake volume, and shorter antibiotics treatment duration were also significantly associated with low IP (**Table 1**). Compared to infants with low IP, neonates with high IP had fewer days of MOM feeding (4 days vs. 5.5 days, P<0.01) and less total MOM volume (123.4 ml/kg vs. 263 ml/kg, P<0.01) as well as longer duration (>3 days) antibiotics use (37.5% vs. 18.5%, P=0.03). We adjusted host factors associated with IP and fit a generalized logistic regression model. Newborns who were fed MOM for ≥4 days during the first week were demonstrated to be 10.3-fold more likely to have low IP than those who were fed MOM for <4 days [adjusted odds ratio (aOR): 10.3, 95% CI: 3.21-33.33] (**Table 2**). Additionally, newborns who had longer antibiotics treatment (≥3 days) were 2.6 times more likely to have high IP, however this association was mitigated when adjusting for confounders like PMA. This result is in line with our previous observations that antibiotic use is significantly more common in the early GA subjects (92% in <28 weeks GA *versus* 32% in >28 weeks GA, P<0.001) (16). Statistical dependence analyses showed that the cumulative intake volume of MOM prior to the IP measurement was the most significant factor associated with IP (P <0.001, q value <0.01, HSIC statistic=1.53 and 1.46), at a significance level even higher than host factors including GA (P <0.001, q value <0.01, HSIC statistics=1.12), PMA (P = 0.01, q value = 0.015, HSIC statistics=0.93), and body weight (P = 0.01, q value = 0.035, HSIC statistics=1.12) (**Table S2**).

**TABLE 2.** Odds ratio for factors associated with Low IP Adjusted for postmenstrual age (PMA) and birth weight (BW)

|  | OR | 95% CI | p-value<sup>d</sup> | Adjusted OR<sup>c</sup> | 95% CI | p-value<sup>d</sup> |
| --- | --- | --- | --- | --- | --- | --- |
| Duration of antibiotics use<sup>b</sup> |  |  |  |  |  |  |
| ≤ 3 days | 2.65 | 1.12, 6.25 | 0.02 | 1.56 | 0.58, 4.16 | 0.37 |
| > 3 days | 1.0 (Ref) |  |  | 1.0 (Ref) |  |  |
| Duration of MOM feeding<sup>b</sup> |  |  |  |  |  |  |
| ≥ 4 days | 7.04 | 2.5, 19.6 | <0.01 | 10.30 | 3.21, 33.33 | <0.01 |
| < 4 days | 1.0 (Ref) |  |  | 1.0 (Ref) |  |  |

<sup>a</sup>Fisher's exact test was used to calculate p value for categorical variable. Student's t-test was used for continuous variables (BW, GA, PMA, APGAR score at 1 minute and 5 minutes). IP was calculated as the ratio of urine Lactulose (La) and Rhamnose (Rh) and La/ Rh < 0.05 was defined as low IP.
<sup>b</sup>Variable measured during the time-period starting from enrollment day (within 1 to 4 days after birth depending on clinical stability) until the day when IP was measured (day 8±2 post-birth).
<sup>c</sup>Adjusted OR model includes PMA and BW.
<sup>d</sup>p-value calculated using logistic regression.
**ABBR:** IP, Intestinal Permeability; PPRM, Premature Preterm Rupture of Membranes; BW: Birth Weight; VLBW, Very Low Birth Weight; MOM, Mothers own breastmilk; GA, Gestational Age; PMA, Postmenstrual Age; OR, Odds ratio; CI, confidence interval

### Breastmilk intake is associated with improved intestinal barrier integrity

Unfortunately, mothers who deliver preterm often produce less milk than those who deliver term, and milk administration is often delayed especially in early preterm infants (27). Formula and/or pasteurized donor human breastmilk (PDHM) is often a necessary dietary supplement. Only 55.7% of neonates in the cohort were exclusively breastfed (N=63), others had either complemented with formula (N=31), or PDHM (N=12), or were fed exclusively formula (N=9) (**Fig. 2A**). For this reason, we investigated IP in neonates grouped by feeding types. Exclusive formula feeding was significantly associate with high IP, either in number of days (P=0.02) or the intake volume (P=0.03, **Table 1**). However, when formula was used in combination with MOM, even at a minor portion (35.2%±31.7%, mean±s.e.), IP was significantly lowered to a level that is no different than exclusive MOM (**Fig. 2B**). Infants whose diet was supplemented with PDHM in addition to MOM had similar IP to the exclusive MOM group. We further investigated how much MOM is “sufficient” relating to improved IP during the first week post-birth. A highly elevated IP was observed in infants who received no MOM (exclusive formula or no feed), and a rapid decrease in IP was inversely correlated with increased MOM intake volume (**Fig. 2C**). Discriminatory machine learning schemes suggested that a threshold around 150-180 ml/kg of cumulative intake of MOM by 7-10 days of age is associated with low IP. Together our results indicate that sufficient MOM, used alone or combined with other forms of feeding, significantly impacts IP in early preterm infants. Even more importantly, these results imply that the benefits of breastmilk feeding are beyond the nutrition alone but extend to postnatal intestinal barrier maturation.

**FIG 2.**
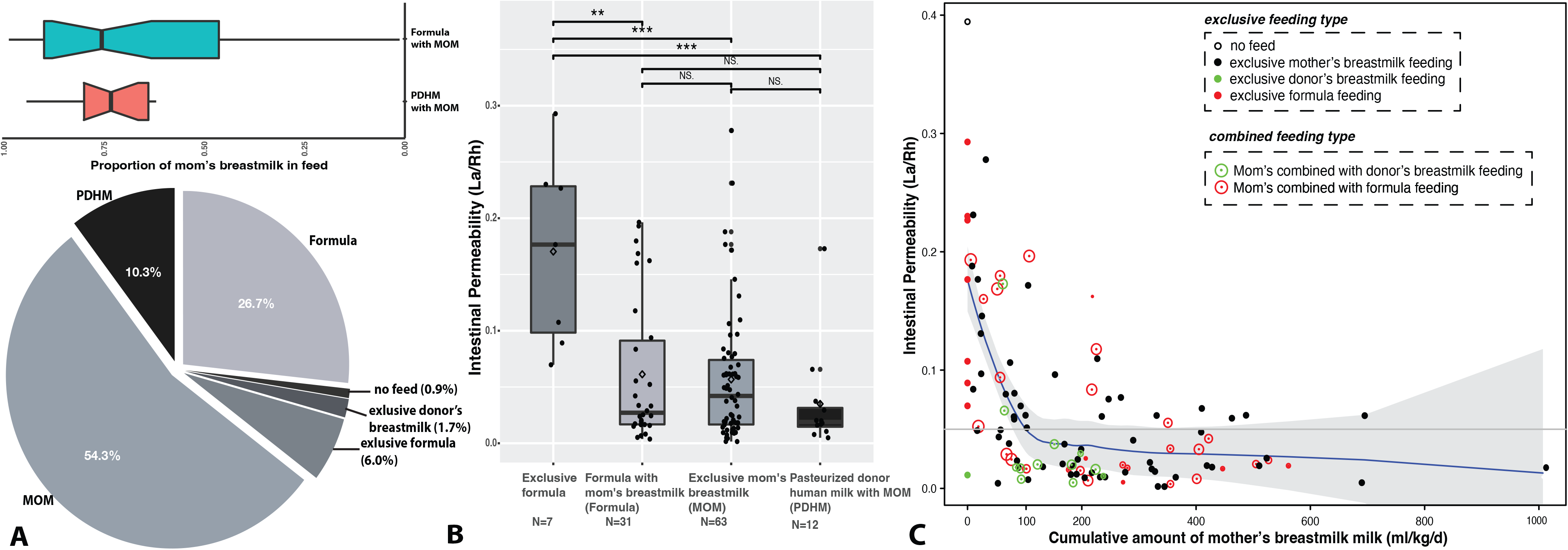
Pie chart of feeding types of the preterm infant population in this study (A). Abbr: MOM: mother’s breastmilk feeding; PHDM: pasteurized human donor’s milk. Boxplot of the IP measurement grouped by feeding types (B). Correlation between intestinal permeability and the cumulative amount of mom’s own breastmilk feeding (ml/kg) for a total of 113 enrolled preterm infants 24^0/7^-32^6/7^ weeks of gestation were enrolled (C). IP was calculated using the ratio of urine Lactulose (La) and Rhamnose (Rh), low and high IP was defined by a La/Rh >0.05 or ≤0.05, respectively. The total amount of mom’s own breastmilk feeding was calculated as sum of the daily amount of milk intake per kilogram bodyweight until d7-10 when the IP was measured. Initial feed was calculated based on 10 ml/kg expressed breast milk between the first and fourth day of life depending on clinical stability. After 3-5 days initial feeds, feedings were advanced by 20 ml/kg/d until 100 ml/kg/d was reached. Plotted are interquartile ranges (IQRs, boxes), medians (line in box), and mean (diamond). Significance value was calculated using Wilcoxon rank sum test. Star sign (*) denotes the level of significance. “NS” denotes non-significant.

### Increased *Bifidobacterium* species abundance correlates with improved intestinal barrier integrity

We further performed high-resolution characterization of intestinal microbiota in 517 fecal samples, using both short-read sequencing of the V3V4 region of the 16S rRNA gene on an Illumina HiSeq 2500 instrument (300PE, N=472), and long-read sequencing of the full-length 16S rRNA gene on PacBio Sequel II platform (N=192). For short-read sequencing, we obtained a total of 25,838,078 high-quality, non-chimeric ASVs (Amplicon Sequence Variants) after the assembly of forward and reverse reads and quality assessment, representing 51,165±620 (mean±s.e.) ASVs per sample (**Table B** at https://doi.org/10.6084/m9.figshare.19723252.v1). On the other hand, long-read sequencing generated using the Circular Consensus Sequences (CCS) yielded 1,271,873 high-quality full-length 16S rRNA sequences or 992.9±16.8 (mean±s.e.) non-chimeric ASVs per sample. The full-length 16S rRNA gene sequences (1,462 bp on average) extended the partial V3V4 region (428 bp on average) 3.2 times, and afforded species level assignment for 87.6% of the long-read ASVs (remaining were not assigned due to a lack of reference), compared to 15.3% for the short-read ones (**Table D** at https://doi.org/10.6084/m9.figshare.19723252.v1**, Fig. S2**). Using samples sequenced by both methods, taxonomic assignments for long-read ASVs were conveyed to short-read ASVs using perfect sequence match, thus achieving species assignment in 65.3% of short-read sequences (**Table E** at https://doi.org/10.6084/m9.figshare.19723252.v1).

In total 508 ASVs belonging to 212 species in 15 orders and 6 phyla were identified (**Table A-C** at https://doi.org/10.6084/m9.figshare.19723252.v1). The four most abundant taxa were *Klebsiella pneumoniae*, *Escherichia coli*, *Staphylococcus epidermidis*, and *Enterobacter* spp. These taxa were predominant (>50% relative abundance) and dictated four distinct community types (**Fig. S3**). These four taxa belong to two classes Enterobacteria (*K. pneumoniae*, *E. coli*, and *Enterobacter* spp.) and Bacilli (*S. epidermidis*) and were highly prevalent (present in 86.2-94.8% samples) in both high and low IP subjects (**Fig. 3A**). They are also known “first colonizers” of the infant gut (15, 28, 29). Five other taxa, including *Enterococcus faecalis, Clostridium perfringens, Proteus mirabilis, Bifidobacterium breve,* and *Veillonella dispar*, were found to contribute to 17.4% of all sequences and detected in 47.7-86.6% of all samples. These obligate and facultative anaerobes were considered the “succession” microorganisms that succeed to the first colonizers (15, 30–32). Together these nine taxa accounted for 76.0% of all sequences in this dataset. Remaining sequences were from a diverse array of obligate and facultative anaerobes (**Fig. S3** cluster 5).

**FIG 3.**
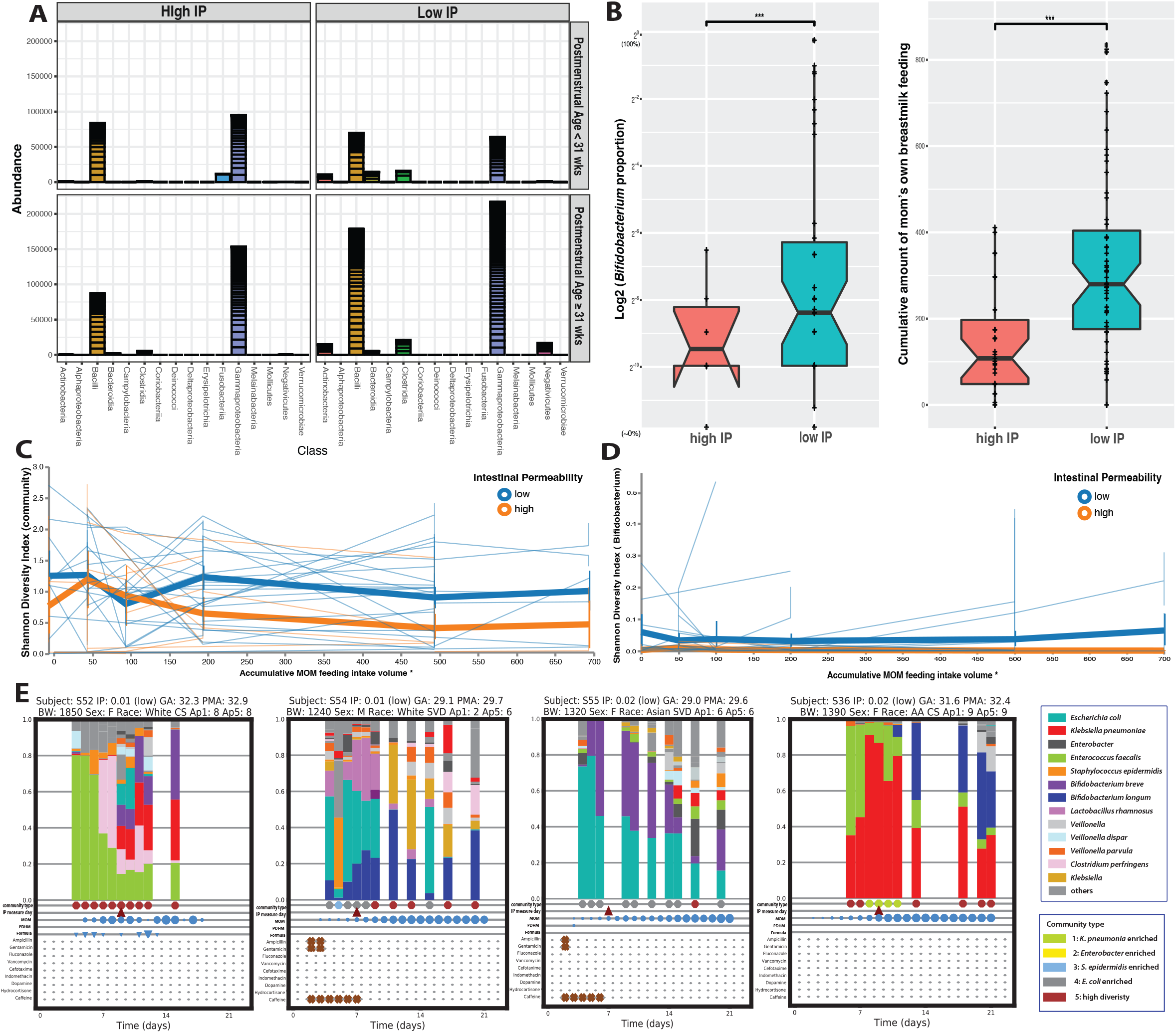
Microbial biomarkers and breastmilk feeding in early preterm subjects with high and low IP. Abundance of bacterial groups stratified by postmenstrual age at study day 7-10. It indicates the Actinobacteria (*Bifidobacterium*) and Clostridia (Clostridiales) that were mainly observed in low IP subjects but not in high IP subject (red) (A). The abundance values of read count for each ASVs are stacked in order from greatest to least, separate by a horizontal line. Boxplot of the *Bifidobacterium* relative abundance and cumulative amount of mom’s breastmilk feeding (ml/kg) during the first 7-10 postnatal days in subjects with high or low IP (B). IP was calculated using the ratio of urine Lactulose (La) and Rhamnose (Rh), low and high IP defined by a La/Rh >0.05 or ≤0.05, respectively. Plotted are interquartile ranges (IQRs, boxes), medians (line in box), and mean (diamond). Significance value was calculated using Wilcoxon rank sum test. Star sign (*) denotes the level of significance. “NS” denotes non-significant. Volatility plot to demonstrate the fluctuation of microbial community diversity (C) (characterized as Shannon diversity index) and *Bifidobacterium* diversity over MOM feeding volume in high or low IP groups (D). Plot was generated in QIIME (2019.10 vers) (106). Non- overlapping of the vertical error bar at each measuring point was considered significantly different. Temporal characterization of intestinal microbiota of early preterm infants to profile changes over the first 21 days post-birth (E). Taxonomic profile was generated using 16S rRNA gene sequencing. Community type is shown in Fig. S3 heatmap clusters. The dates when IP was measured, MOM, PHDM, formula feeding day, antibiotics administration are shown in the plot. Each circle is sized proportionally the feeding volume. Abbr: MOM: mother’s own breastmilk feeding; PHDM: pasteurized human donor’s milk.

A zero-inflated negative binomial random effects model (ZINBRE) was applied to investigate microbial biomarkers correlated with IP. *B. breve* was the taxa the most significantly associated with low IP (P < 0.001) during the first 7-10 days after birth (**Table S3B,** **Fig. 3B, S4B**). The low IP group had significantly higher levels of *B. breve*, more *Bifidobacterium* overall, and more MOM. An adaptive spline logistic regression model was used independently to confirm the association between *B. breve* to IP and MOM (**Fig. S4C,D**). Other phylotypes associated with MOM or PMA were shown in **Table S3.** The high IP-associated ASVs of *S. epidermidis*, *E. coli*, *Parabacterioides distasonis* were associated with early PMA (**Table S3A**). *Veillonella dispar* was revealed to strongly associate with later PMA (P<0.001) but not with IP. *S. epidermidis* and *E. coli* were also associated with less MOM during the first week (**Table S3C**). *B. breve* was in 71.7% of samples containing *Bifidobacterium*, followed by *B. longum* (21.7%). The other *Bifidobacterium* species were either rare or in very low abundance (<0.1%). Temporal microbiota profiling indicated that *Bifidobacterium* species reached higher abundance ∼5-20%) after >3d of MOM (**Fig. 3E**, https://doi.org/10.6084/m9.figshare.19709923.v1). When stratified by major feeding types, *Bifidobacterium* was mostly abundant in exclusive MOM or MOM supplemented with formula (**Fig. S4A**). We plotted community diversity against MOM feeding volume in function of time and observed that low IP infants had significantly higher diversity microbiota and higher diversity *Bifidobacterium* species, when MOM reached >150 ml/kg of cumulative intake within the first week (**Fig. 3C-D**). It is worth noting that MOM is a critical but not the only contributor to the abundance of *Bifidobacterium*. 15% of the subjects received no MOM had >1% *Bifidobacterium* and 32.5% had detected level of *Bifidobacterium* (>0.1%). Overall this result further supports the importance of achieving the critical threshold of MOM intake and its critical association with low IP.

### Population dynamics of *Bifidobacterium* species in early postnatal colonization

Phylogenetic analyses of full-length 16S rRNA gene sequences demonstrated that *B. breve* forms a monophyletic clade and the four most abundant ASVs were nearly identical, while *B. longum* was more phylogenetically diverse with four distinct clades (**Fig. 4A,B**). Clade I was the most abundant and represented *B. longum* subsp*. longum*, while *B. longum* in the other three clades II-IV was in low abundance. ASVs assigned to *Bifidobacterium* showed high sequence diversity (**Fig. 4A**) as well as inter- and intra-subject variability (**Fig. 4C**), in that multiple ASVs can be detected in the same subject and a single ASV can be detected in multiple subjects at multiple time points. For instance, 35 *B. longum* ASVs of four different clades were observed in one subject. Further, some ASVs (i.e., unclassified *Bifidobacterium* spp.) were only observed in infants with early PMA (<33 weeks) while others did not vary in abundance across PMA (i.e., *B. breve*), supporting a high subspecies-level diversity and population dynamics in preterm infant gut community.

**FIG 4.**
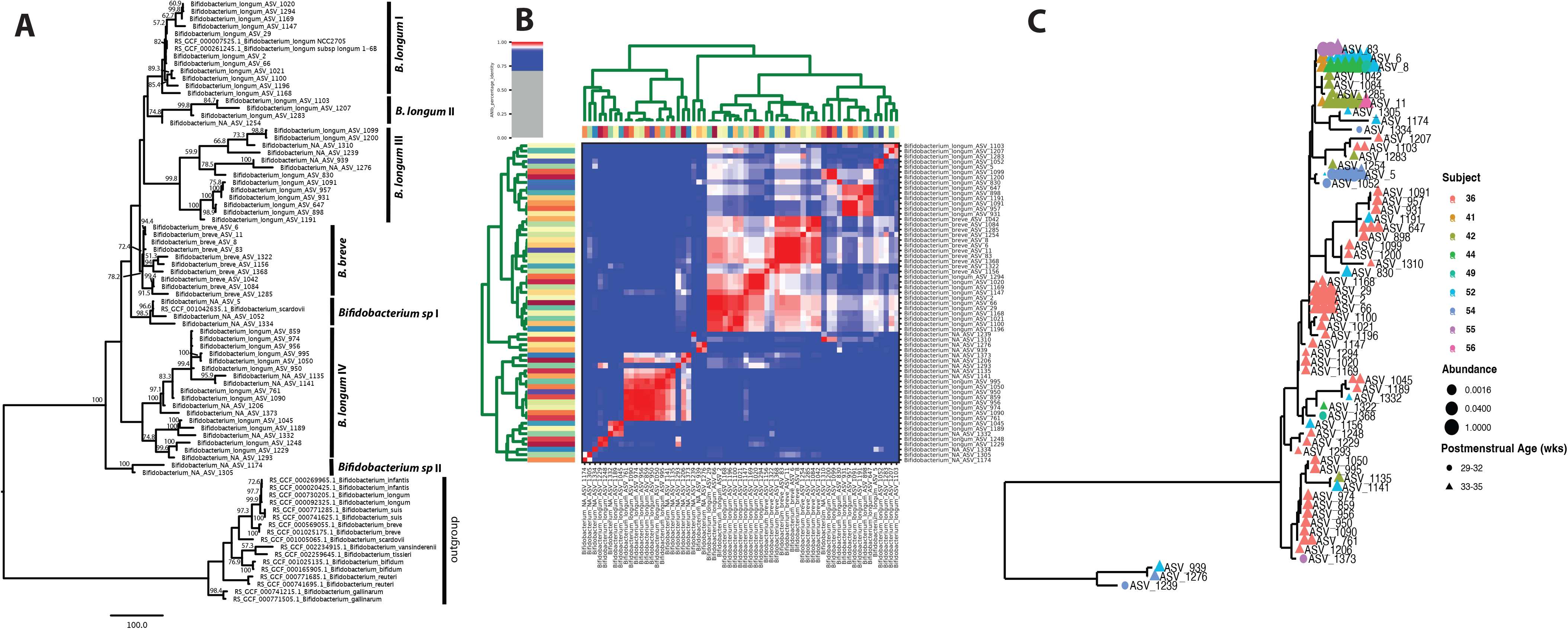
Phylogenetic tree constructed using 81 unique, full-length 16S rRNA gene ASV sequences of *Bifidobacterium* (A). ANI clustering of full-length 16S rRNA gene sequences (B). Phylogenetic tree of *Bifidobacterium* ASVs in stool microbiota of cohort (C). All full-length 16S rRNA genes assigned to *Bifidobacterium* were used in the analyses. Color denotes individual subjects.

To characterize the genome content of *Bifidobacterium* species, we performed whole metagenomic sequencing of 30 samples with >10% *Bifidobacterium* species using an Illumina NovaSeq 6000 platform (**Table A** at https://doi.org/10.6084/m9.figshare.19723255.v1) and generated 26 *B. breve* and four *B. longum* nearly complete metagenomic-assembled genomes (MAGs) (**Table B** at https://doi.org/10.6084/m9.figshare.19723255.v1). We further performed metagenomic sequencing of two samples using Pacific Bioscience Sequel II platform, which afforded one closed and one nearly complete genomes of *B. breve* strains. The closed genome was 2.34M in size (**Fig. S6, Table C** at https://doi.org/10.6084/m9.figshare.19723255.v1), similar to the median *B. breve* genome size of 2.33M on NCBI. For pangenome analysis, we supplemented the 26 *B. breve in-house* MAGs with 107 published genomes (**Table A** at https://doi.org/10.6084/m9.figshare.19709917.v2) and the four *B. longum* MAGs with 310 published genomes (**Table B** at https://doi.org/10.6084/m9.figshare.19709917.v2) to identify homologous gene clusters (HGCs) (**Tables C-D** at https://doi.org/10.6084/m9.figshare.19709917.v2). Among the total of 4,922 *B. breve* HGCs, 54.2% were considered dispensable (present in <10% genomes), 29.4% were core (present in >95% genomes) and the rest were accessory (**Table E** at https://doi.org/10.6084/m9.figshare.19709917.v2). The pangenome of *B. longum* (7,265 HGCs) was roughly twice the size of *B. breve* (3,363 HCGs), although the two species core genomes were similar (1,511 vs. 1,448 HCGs). The large pangenome size of *B. longum* may reflect its broader host range that includes both infant and adult intestines than *B. breve* or *B. infantis,* which were exclusively observed in infant gut (33). In particular, the genes involved in the fructose 6-phosphate phosphoketolase- dependent glycolytic pathway for ATP-efficient carbohydrate catabolism, or “bifid shunt”, are conserved in both species (**Fig. S7**). Further, *B. longum*’s dispensable genome, which comprised 46.3% of its pangenome (2,666 HGCs), was smaller than that of *B. breve* (54.2%, 3,363 HCGs) in both size and proportion, indicating a high genome plasticity in *B. breve*.

We identified 46 genes specific to *B. breve* colonizing infants with low IP (**Table F** at https://doi.org/10.6084/m9.figshare.19709917.v2). While a large number of these genes have unknown functions, others encoded functions such as glycosyl transferases, glycosyl hydrolases, cell surface adhesion and transport, polysaccharide biosynthesis, quorum sensing, and phage integration. Further, a number of functions were significantly enriched in these genomes compared to the species’ genomes publicly available (adjusted q-value < 0.05, **Table F-I** at https://doi.org/10.6084/m9.figshare.19709917.v2), such as cation transmembrane transporter activity, glucuronate isomerase, methyladenine glycosylase, glycosyl hydrolase family 59, 2, 85, 30, bacterial rhamnosidase A and B. Of note, *B. breve* HGC profiles appears to be highly similar within subjects, indicating that *B. breve* genomes detected at different time points in the same infants shared greater similarity than those from different subjects (**Fig. S7**, **Table J** at https://doi.org/10.6084/m9.figshare.19709917.v2). Together, compared to *B. longum, B. breve* colonizing infants with low IP has a high genome plasticity and enriched genetic features in carbohydrate metabolism and transportation that underlies the species strong niche adaptive capabilities.

### Specialized human milk oligosaccharides assimilation capabilities of Bifidobacterium strains in early preterm infants

As both *Bifidobacterium* species abundance and MOM were associated with postnatal intestinal barrier maturation, we next investigated whether these two factors were linked through the ability of *Bifidobacterium* species to utilize the oligosaccharides present in breastmilk. Previously characterized major HMO utilizers like *Bacteroides* species and *Lactobacillus* (34, 35) were largely absent from our cohort (https://doi.org/10.6084/m9.figshare.19723252.v1), indicating that *Bifidobacterium* species likely provide the genetic capabilities to metabolize HMOs. We thus examined the set of genes encoding extracellular hydrolases, sugar transporters, and intracellular hydrolases (**Table S4**), which comprise the machinery necessary to uptake and metabolize HMO substrates to feed the central fermentative metabolism (36–38).

Intracellular HMO utilization functions were exclusively found encoded by both *B. breve* and *B. longum*. We examined eight essential extracellular enzymes and their homologs (details in methods section) known to be required in extracellular breakdown of HMOs into smaller molecules that are then transported intracellularly. Interestingly, none of these extracellular enzymes were found in this cohort. We investigated five essential bacterial ABC transporters and homologs involved in the import of various oligosaccharides, known to have a high specificity for HMOs conferred by substrate-binding protein (SBPs) domains (39). Both *B. breve* and *B. longum* contained *gltA* (**Table S4A**), a gene considered crucial to the import of lacto-N-tetraose (LNT). LNT comprise the core HMO structure that is catabolized via lacto-*N*-biose (LNB) intermediates (40). Further, a family 1 solute binding proteins (F1SBP) gene cluster Blon_2177, was found in both *B. breve* and *B. longum* (**Table S4B**). This cluster was found critical in the import of non-fucosylated type 1 oligosaccharides (41). None of the *B. longum* strains but the majority *B. breve* strains of this cohort (92.4%) harbor the LNnT (lacto-N- neotetraose) transporter that is encoded by *nahS*. These findings indicate both *B. breve* and *B. longum* could transport LNB and LNT, while *B. breve* can further metabolize LNnT.

We then evaluated the capability of consuming the transported oligosaccharides, and, compared to *B. longum,* we revealed expanded metabolic capabilities of *B. breve* of this cohort to utilize a variety of HMO molecules including fucosylated or sialylated forms, in addition to the neutral types of HMOs (*i.e.*, LNB, LNT, LNnT). 17 key glycoside hydrolases (GH) involved in essential HMO degradation and utilization were investigated (**Table S4C**). Key intracellular enzymes GH2 (β-1,4-galactosidases, LacZ2/6), GH112 (GNB/LNB phosphorylase, *lnpA*), GH20 (β-N-acetylglucosaminidase), and GH42 (β-1,3-galactosidase, *lntA*, bga42A) are highly conserved in both *B. breve* and *B. longum*. These enzymes lack transmembrane domains or signal peptide sequence and are required to degrade HMOs intracellularly (42). While almost all *B. breve* contained GH95 α-fucosidase (*afcA*, homolog to Blon_2335), GH33 α-sialidase (homolog to Blon_0646), and GH20 β-N-acetylglucosaminidase (*nahA*, homolog to Blon_0459) (**Table S4C**), only a small portion of *B. longum* (∼10%) contained these enzymes. Further, *B. breve* present in these preterm infants carries gene encoding GH29 α-fucosidases more often (53.8% vs. 12.7%) than *B. breve* isolated from other sources obtained from GenBank. The presence of GH29 α-fucosidase genes underlines the capability to consume fucosylated oligosaccharides such as 2’-fucosyllactose (2’-FL) and larger fucosylated HMOs such as lacto-N-fucopentaose (38, 42). The GH29 containing *B. breve* strains in our cohort also encode GH95. In fact, GH29 and GH95 α- fucosidases are highly complementary since they target specific substrate of α-1,3/4 and α-1,2 fucosyl linkages, respectively (42), and the activation of both enzymes enables degradation and utilization of a higher variety of HMOs. Moreover, a prominent gene cluster termed FHMO (Fucosylated Human Milk Oligosaccharide) that contains both GH29 and GH95 α-fucosidases coding genes was observed in some *B. breve* strains but is largely absent from *B. longum* (**Table S4D**). This cluster was reported to enable *B. breve* strains to preferentially consume fucosylated HMOs over neutral HMOs during early bacterial growth (42). In particular, the putative fucosyl lactose SBP (BLNG_1257) present in this cluster confers glycan binding specificity and is consistently present in *B. breve* strains of this cohort but rarely in other *B. breve* in GenBank. Overall, our results revealed an expanded, specialized HMOs assimilation capability of *B. breve* strains, conferring a competitive growth advantage in the gut of this preterm infant cohort when fed breastmilk.

### Host-derived glycoproteins utilization is limited to B. breve in early preterm infants

Besides HMOs, the host-derived glycoproteins such as mucin and proteoglycan (mucus or milk) are critical carbon sources to bacteria in the infant intestinal microenvironment. Human glycoproteins are often heavily sulfated and could not be metabolized without bacterial glycosidases (43, 44). We investigated two sulfatase-encoding gene clusters essential in sulfatase metabolism *ats1* and *ats2* (45, 46), and they each encode glycosulfatases and accompanying anaerobic sulfatase- maturing enzymes (anSMEs) with an associated transport system and transcriptional regulator (46). The primary mucin degradation capabilities in this cohort are shown to be limited to *B. breve* strains (**Table S4F**), as the two clusters are present in 100% of *B. breve* in our cohort and ∼70% of all *B. breve* genomes available. *B. longum* rarely harbor *ats1* and no strains carry *ats2*.

In addition to sulfated residues, more than half of human colonic mucin oligosaccharides also contain sialic acid residues (47). The release of sialic acid is an initial step in the sequential degradation of mucins and sialylated HMO substrates (46, 48). Hence, we investigated the two gene clusters essential for the uptake and metabolism of sialic acid, *nagA2-nagB3* cluster (Bbr_1247, Bbr_1248) and the *nan-nag* cluster (Bbr_0160-0172) (49–51). These two gene clusters are highly conserved in *B. breve* while only present in 14% of *B. longum* genomes (**Table S6E**). Our results demonstrate the capability of foraging sulfated and/or sialylated host-derived glycoprotein is attributed to *B. breve* strains in this cohort. This metabolic versatility of *B. breve* may greatly improve its fitness and facilitate its mucosa adherence, hence facilitating the colonization under nutrient- or energy-limited conditions in the preterm infant gut environment.

## DISCUSSION

Early preterm neonates are a vulnerable and challenging population that often requires intensive medical care. As a result of their premature birth, these neonates often have an aberrantly permeable intestinal barrier that fail to limit bacterial translocation. Our group has previously reported positive associations between persistently elevated intestinal permeability and delayed feeding, prolonged antibiotics exposure and altered development of the intestinal microbiota, and a lack of progressive increased abundance of *Clostridiales* (15, 16). These *Clostridiales* became abundant mostly at the end of the second week post-birth, this is after the extensive barrier maturation that occurs during the first week. In this study, we determined the minimal intake of maternal breastmilk necessary to significantly lowered IP, and identified specific *Bifidobacterium* species and strains as the biomarkers associated with low IP development in preterm infants first week of life.

We posited the benefits of breastmilk extend beyond nutrition and include improved gut barrier function, and that the two factors associated with reduced IP, MOM feeding and *Bifidobacterium* strains, at least in part, are linked by the capability of the *Bifidobacterium* to metabolize human milk oligosaccharides (illustrated in **Fig. 5**). To investigate this link, we evaluated the carbohydrate metabolizing capabilities of *Bifidobacterium* strains and uncovered a complement of genes dedicated to utilizing a wide variety of HMO molecules as well as host-derived glycoproteins. These genetic features were enriched in preterm infant gut-associated *Bifidobacterium* strains compared to those isolated from other sources like dairies or adult gut. Our results are concordant with previous studies that the establishment of a bifidobacterial dominant community was facilitated by specific gene clusters supporting HMOs metabolism, which are absent in many adult associated bifidobacterial strains (52–55). The functional characterization of the contribution of *B. breve* metabolizing MOM to low IP would be critical to its translational significance. Future studies modeling both transcriptional activities of bifidobacterial biomarkers and host responses in a longitudinal design is warranted to address the causal-effect relationships of MOM and *Bifidobacterium* on intestinal barrier maturation. Further, the production of short chain fatty acids via carbohydrate consumption by bifidobacteria, particularly acetate and butyrate, was demonstrated to correlate with their anti-inflammatory properties and promoted the defense functions of the epithelium (56–58). Together, our study supports the notion that intestinal barrier function can develop postnatally, and this process could be induced through supplementation of breastmilk substrates as well as *Bifidobacterium* strains that consume them. These elements are promising therapeutic targets to reduce NEC and other life-threatening conditions associated with intestinal hyperpermeability.

**FIG 5.**
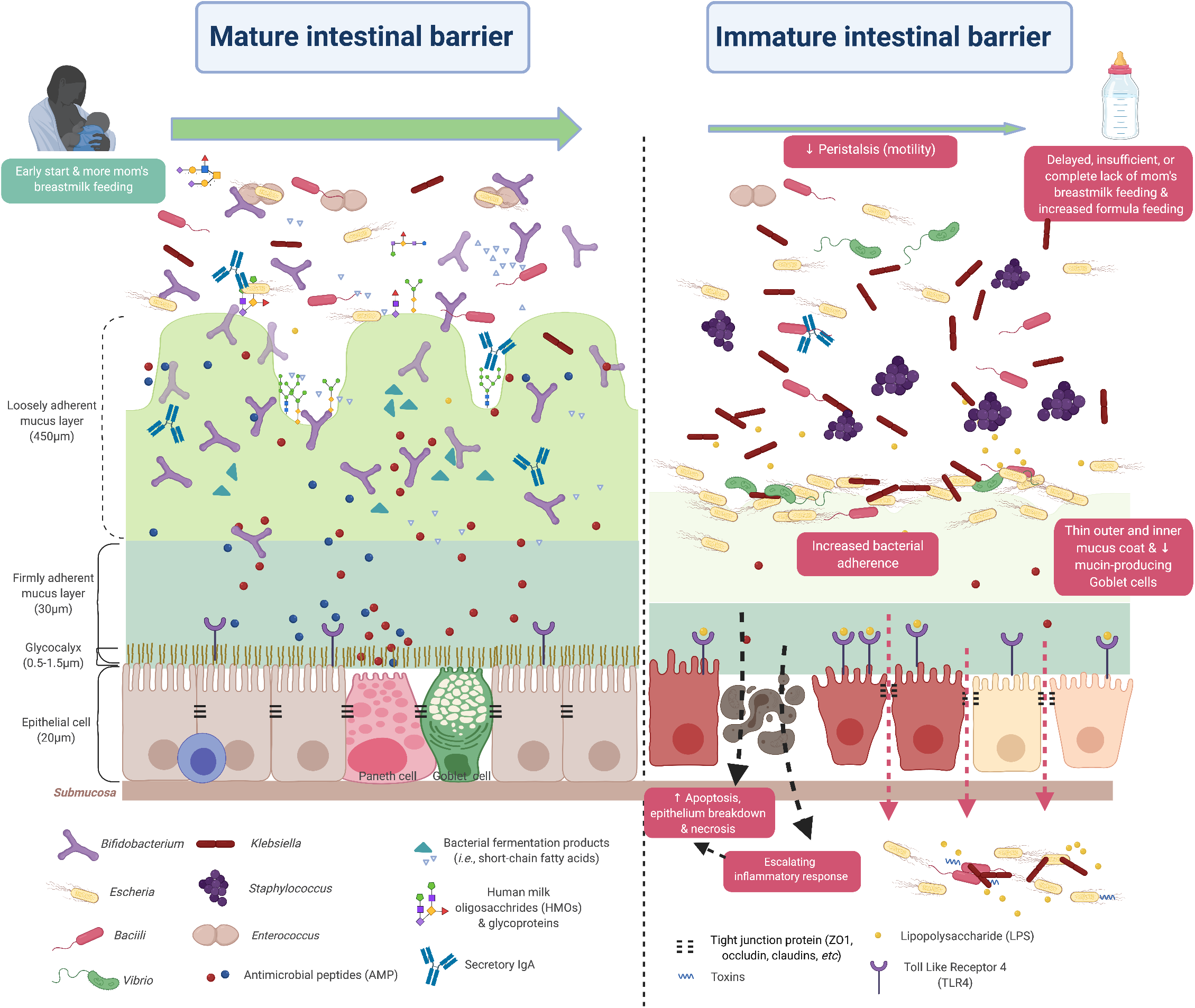
Illustration of the mature and immature intestinal barrier in neonates. Peristalsis (reduced intestinal motility), maldigestion of nutrient sources and a compromised gut barrier may render the mucosa susceptible to invasion by the opportunistic pathogens in gut environment. The resulting imbalance between epithelial cell injury and repair leads to a vicious cycle of maldigestion, bacterial invasion, immune activation and uncontrolled inflammation. Illustration not drawn to scale. Created with BioRender.com.

*B. breve* is a known dominant *Bifidobacterium* species in both preterm and term infant gut microbiota (59) and was also observed in breastmilk and vaginal microbiota (60, 61). In human, *B. breve* appears to be exclusively in these environments and is largely absent in adult gut. The factors contributing to *B. breve* persistence in infants are not well understood. Most studies were performed using the type strain *B. breve* ATCC 15700 (JCM 1192), which has limited ability to consume HMOs (62, 63). As demonstrated by us and others, strains of *B. breve* vary greatly in their capabilities to metabolize HMOs (55). The *B. breve* strains in our cohort displayed extensive enzymatic capability designed to efficiently utilize a broad range of dietary and host-derived carbohydrates and thus maximizing their colonization in the infant intestinal environment. In particular, we demonstrated that LNnT utilization was exclusively limited to strains of *B. breve*. Growth on LNnT was shown *in vitro* to enable *B. infantis* to outcompete other species such as *Bacteroides* (64). LNnT can be fermented by specific strains of *Bifidobacterium* only found in infant gut (65). Digestion of neutral HMOs (*i.e.*, LNT, LNnT) was actually shown to induce a significant shift in the ratio of secreted acetate to lactate compared to the catabolism of the simpler carbohydrates they contain (66). Further, GH29 α-fucosidase, an uncommon enzyme correlated to the ability to grow on fucosylated HMOs (38), was only enriched in *B. breve* strains in this cohort.

The presence of key gene sets expands *B. breve* metabolic capabilities (*i.e.*, FHMO, GH29, GH95), and is reminiscent to those found in *B. infantis* ATCC 15697, the model strain that can also consume a broad repertoire of HMOs (41, 67). Previous clinical trials administrating *B. breve* strains in early preterm infants yield contradicted results, which may relate to the different strains selection. For example, Kitajima and co-authors reported a *B. breve* strain BBG could colonize the immature bowel effectively with significantly fewer abnormal abdominal signs and greater weight gain in VLBW infants (68). However, the clinical trial of the type strain BBG-001 in very preterm infants observed no evidence of benefit in terms of preventing NEC and LOS (69). These data highlight the importance of strain characterization in prophylactic supplementation of live biotherapeutics. Further characterization of these key genes will be necessary to understand the range of oligosaccharides *B. breve* strain can transport and consume. Strains collection of *B. breve* isolated from both preterm infants with rapidly lowering IP and healthy term infants should be established to achieve this important goal.

The specialized HMOs and glycoprotein utilization capabilities of *B. breve*, particularly the sulfated and sialic residues degradation, further confers a competitive capability that improve *B. breve* fitness and facilitate its adherence and colonization of the gut mucosa (70). The release of sialic acid is an initial step in the sequential degradation of mucins and sialylated HMO substrates (46, 48), and the ability to utilize the heavily sulfated mucin glycoprotein and sialic residues were found to be highly correlated (46, 49). Sialic acid concentrations are highest in colostrum in preterm infants but decrease by almost 80% after 3 months (71). Further, breastmilk from mom who delivered preterm was reported to be a rich source of oligosaccharide-bound sialic acids, with 20% more sialic acid residues than breastmilk from term mothers and 25% more than that found in formula (72). A recent *in vivo* study showed that sialylated HMOs are on the causal pathway of a microbiota-dependent infant growth outcome, hence were considered the most growth-discriminatory HMO structures (73). Interestingly, and supporting its importance in infant health, only strains of *B. infantis* and *B. breve* isolated from infant gut have been reported to be capable of utilizing sialic acid and sialylated lacto-N-tetraose as sole carbon source (54, 74, 75). A few *B. breve* strains were actually reported to preferentially consume sialylated HMOs, in particular sialyl- LNT b (LSTb), sialyl-lacto-N-hexaose (S-LNH) over neutral HMOs (38, 49). Given that bacteria with pathogenic potentials are capable of utilizing sialic acid, *B. breve* strains could rapidly sequester sialic acid away from these pathogens and offer nutritional immunity, *i.e.,* sequester nutrients to limit infection, thus contributing to a healthy intestinal environment (76). It would be highly insightful to further characterize maternal HMOs variations in MOM and the composition of specific formula in addition to the information of HMOs assimilation capability of bifidobacterial strains, for comprehensive understanding of the essential factors attributed to postnatal intestinal maturation.

HMO utilization by *Bifidobacterium* species in this cohort appears to be exclusively an intracellular process, which would unlikely allow for cross-feeding of intermediates with other gut bacterial species. Extracellular digestion of HMOs would afford fucose and sialic acid monomers to be cross-fed to other bacteria, some of which with pathogenic properties (77). *Bacteroides spp* that are largely absent in this cohort are known to employ exclusively extracellular process in HMO utilization (64). The “internalize, then degrade” approach for HMO consumption is a critical *Bifidobacterium* property that affords protection against infection for the infants. Interestingly, the preference for intracellular digestion of HMOs is not conserved across all infant gut *Bifidobacterium* species or strains. A recent study revealed *Bifidobacterium* in the gut microbiome of breast-fed Malawi and Venezuela infants similarly employed an intracellular HMO digestion strategy, while *Bifidobacterium* in a cohort of US infants fed formula and breastmilk preferentially employed extracellular HMO digestion strategies (36). The difference may relate to galacto-oligosaccharides (GOS) transporter genes present in strains that internalize HMOs to metabolize them, especially the GNB/LNB- BP (*GltA*) gene (36, 78), though the mechanisms remain unclear.

Our study highlights the strong potential for the prophylactic administration of specific *B. breve* strains early in life along with specific HMOs to enhance intestinal barrier in preterm neonates. We previously defined a “window of opportunity” of day 8±2 post-birth, for intervention prior to the onset of leaky gut-associated conditions such as NEC (15, 16). Our study proposed the role of breastmilk feeding in promoting the growth of beneficial *Bifidobacterium* species and strains that could consume breastmilk HMOs during that critical window period of time. In the absence of these prophylactic *Bifidobacterium*, the benefit of breastmilk feeding is expected to be dramatically reduced. Counting on the vertical transmission of these *Bifidobacterium* strains from the mothers’ gut or vaginal microbiota, or breastmilk is not reliable and could leave many infants unprotected (79, 80). It is thus critical to gain further mechanistic insight into bifidobacterial-rich microbiota formation in the infant gut by prophylactic supplementation of live biotherapeutics that possess the ability to effectively utilize them. Such understanding will inform the design of clinical interventions with supplementation of HMOs and *Bifidobacterium* as live biotherapeutics prophylaxis to enhance intestinal barrier integrity early in life, and ultimately reduce risk for NEC.

## MATERIALS AND METHODS

### Study cohort and feeding protocol

The study protocol was approved by the institutional review boards of the University of Maryland, Baltimore and Mercy Medical Center. Written informed parental consent was obtained. Eligibility criteria were described previously (16). 113 eligible preterm infants 24^0/7^-32^6/7^ weeks of gestation were enrolled within 4 days after birth from combining cohorts enrolled during June 2013-October 2014 and October 2018-Nov 2019. Prior to study procedures, a complete physical exam including vital signs, weight, height, and head circumference was performed. Demographic, obstetric and clinical, medication exposures, feeding practices and adverse events data were collected from the medical record.

Enteral feeds by the orogastric or nasogastric route were initiated between the first and fourth day of life depending on clinical stability. After initial feeds of 10 ml/kg expressed breast milk or 20 kcal/oz preterm formula daily for 3-5 days, feedings were advanced by 20 ml/kg/d until 100 ml/kg/d was reached. Subsequently, caloric density was advanced to 24 kcal/oz prior to increasing feeding volume by 20 ml/kg/d to 150 ml/kg/d. The total volume of each source of feeds was calculated as sum of the daily amount of milk intake per kilogram of the administered expressed mom’s breastmilk, donor milk or preterm formula from initial feed day till postnatal day 7-10 when the IP was measured. Feedings were held or discontinued for signs of feeding intolerance such as abdominal distension, gastric residuals, or hematochezia, or for clinical deterioration. Pooled pasteurized human donor breastmilk (PHDB) was purchased from Prolacta Biosciences (Duarte, CA, US). PHDB was collected from mothers of term infants who have breastfed for at least 6 months (81).

### *In vivo* intestinal permeability (IP) measurement

In our previous pilot studies that employed a small cohort of neonates (N=37) (15, 16) with IP measured at study day 1, 8±2 and 15±2. It was shown that IP is high within 4 days of birth in all preterm infants with a rapid maturation of the intestinal barrier over the first week of life. Persistently high IP and/or late increase in IP indicate the physiological immaturity of the intestinal tract barrier function. Hence the first 7-10 days in preterm infants is a critical observation period for monitoring IP. Eligible preterm infants received 1 ml/kg of the non-metabolized sugar probes on postnatal day 7-10, which included lactulose (La, Cumberland Pharmaceuticals, Nashville, TN) that is the marker of intestinal paracellular transport and rhamnose (Rh, Saccharides, Inc., Calgary, Alberta, Canada) that is the marker of intestinal transcellular transport. One ml of 8.6 g La +140 mg Rh/100 mL solution was administered enterally by nipple or by gavage via a clinically indicated orogastric tube (82). A minimum of 2 mL of urine was collected over a 4-hour period following administration of the La/Rh dose as previously described (16). La and Rh concentrations were measured by high-pressure liquid chromatography (HPLC) at the University of Calgary (Calgary, Canada). High or low intestinal permeability was defined by a La/Rh >0.05 or ≤0.05 respectively, as validated and applied previously (16). Postmenstrual age at sugar probe dosing was calculated as gestational age at birth plus postnatal age at dosing day (83).

### Fecal specimen collection and nucleic acid extraction

Fecal samples (∼1g) collected daily from enrollment until postnatal day 21 or NICU discharge were stored immediately in 1 ml of DNA/RNA Shield (Zymo Research, Irving, CA, USA). Stool specimen were collected from within the stool mass from the diaper as much as feasible to avoid frequent air exposure. The stool sitting time was 0-3 hours and was collected during diaper change every 3 hours. Urine and fecal samples were archived at -80°C until processed.

Genomic DNA was extracted from homogenized fecal samples using the MagAttract PowerMicrobiome DNA/RNA kit (Qiagen) implemented on a Hamilton STAR robotic platform and after a bead-beating step on a TissueLyzer II (Qiagen) in 96-deep well plates at the Microbiome Service Laboratory (MSL) at the University of Maryland Baltimore (Baltimore, MD, USA). DNA purification from lysates was done on a QIAsymphony automated platform.

### Short-read sequencing of 16S rRNA gene amplicon and whole community metagenomes

PCR amplification of the 16S rRNA gene V3-V4 hypervariable region was performed using dual-barcoded universal primers 318F and 806R as previously described (84). In brief, amplicon pools were prepared for sequencing with AMPure XT beads (Beckman Coulter Genomics, Danvers, MA) and the size and quantity of the amplicon library were assessed on the LabChip GX (Perkin Elmer, Waltham, MA) and with the Library Quantification Kit for Illumina (Kapa Biosciences, Woburn, MA), respectively. PhiX Control library (v3) (Illumina, San Diego, CA) was combined with the amplicon library. High-throughput sequencing of the amplicons was performed on an Illumina MiSeq platform using the 300 bp paired-end protocol. Sequence libraries were prepared from the extracted DNA using the Nextera DNA Flex kit (Illumina; San Dieago, CA) according to manufacturer’s specifications. Libraries were then pooled together in equimolar proportions and sequenced on a single Illumina NovaSeq 6000 S2 flow cell providing an average of 6.5 million pairs of 150 bp reads per library at the Genomic Resource Center at the University of Maryland School of Medicine.

### Long-read sequencing of full-length 16S rRNA gene and whole community metagenomes on Pacific Biosciences Sequel II platform

Amplification of full-length 16S rRNA gene was performed using a dual-barcode, two-step PCR on diluted (1:10) genomic DNA. The first round of PCR amplification of the 16S rRNA full-length gene was performed using universal primers 27F (AGRGTTYGATYMTGGCTCAG) and 1492R (RGYTACCTTGTTACGACTT) following Pacific Biosciences (Menlo Park, CA, USA) specifications for 20 cycles. The cycling conditions for the first-step PCR were 95C for 30sec, 57C for 30sec, and 72C for 60sec. The PCR reaction was then diluted in water (1:5) and amplified with Pacific Biosciences universal forward/reverse 96-plate primers for an additional 20 cycles following Pacific Biosciences specifications. Cycling conditions are as described in manufacture protocol (85). DNA quantification was carried out using the Quant-iT PicoGreen double-stranded DNA assay (Invitrogen) and visualized on a 2% agarose E-gel. The amplicon libraries were normalized and cleaned and concentrated using AmPure XP SPRI beads (Beckman Coulter, Brea, CA, USA) at 0.6X the reaction volume.

Library pools were prepared with SMRTBell Template Prep Kit 1.0 with barcoded adaptors. Libraries were then size-selected on a BluePippen (Sage Science, Beverly, MA) with a cutoff of 5 kb. Sequencing was performed on the Sequel II Platform (PacBio, Menlo Park, CA) with a loading at 60pM. Multiplexed samples were sequenced on PacBio Sequel II cells using the S/P3-C1/5.0-8M sequencing chemistry. Demultiplexing was done with *lima* (version 1.9.0) using default parameters except minimum barcode score 26 and min length 50 bp, both tools are part of the SMRTLink 6.0.1 software package with updated CCS version 3.4.1 (Pacific Biosciences, 2019). Raw reads were assembled via Canu v1.8 and the “-pacbio-raw” protocol (86). Resulting contigs were taxonomically annotated using BLASTN v 2.8.1 (87) and the non-redundant nucleotide database (updated 2019/05/03) to pool all contigs identified under the same species name to form metagenomic bins. Binned contigs were circularized and rotated using “Simple-circularise” (88) and retained if the circularized contigs is in the range of the full genome size according to published closed genomes of that species based on genBank genome database. Metagenome bins were further confirmed using GTDB-Tk v1.1.0 (89). Genomes were annotated using PROKKA v1.13 (90).

### Epidemiological analyses

Covariates identified based on previous literature and biological plausibility were collected at the time of enrollment of the participants and evaluated. Categorical data were compared using Fisher’s exact test and continuous data using Student’s t-test. Multicollinearity between covariates was assessed using Variance Inflation Factor (VIF) and Tolerance, where covariates with VIF >10 were considered collinear. Covariates with p-value < 0.05 in the bivariate analysis were considered confounding factors and were adjusted in the multivariable analysis as random factors. Generalized logistic regression was used to determine the association between IP category and continuous variables including duration of antibiotics and duration of MOM feeding. Analyses were conducted using SAS version 9.4 software (SAS Institute, Cary, NC), code used in this statistical analysis was deposited at https://github.com/igsbma/IP_microbiome/tree/main/statistical_analyses.

### Bioinformatics analysis of intestinal microbiota

For 16S rRNA V3V4 gene amplicon analysis, raw data was demultiplexed and barcode, adapter and primer sequences were trimmed using tagcleaner v0.16 (91). Quality assessment and sequencing error correction was performed using the software package DADA2 v1.14 (92) and the following parameters: forward reads were truncated at position 240 and the reverse reads at position 210 based on the sequencing quality plot, no ambiguous based and a maximum of 2 expected errors per-read were allowed (93). The quality-trimmed reads were used to infer amplicon sequence variant (ASV) and their relative abundance in each sample after removing chimera. The SILVA database (94) release 132 was used to assigned taxonomy. The following criteria were applied on an ASV: 1) at least 400bp in length for long-read sequencing; 2) was observed in at least two samples; 3) at least 5 counts in at least one sample; 4) not assigned to taxonomic groups of Mitochondria or Chloroplast.

For full-length 16S rRNA gene analyses, CCS reads were generated using the ccs application with minPredictedAccuracy=0.99 and the rest of the parameters were default, including minimum 3 subread passes. Demultiplexing was done with lima (version 1.9.0) with minimum barcode score 26 and min length 50bp, both tools are part of SMRTLink 6.0.1 software package with updated CCS version 3.4.1 (Pacific Biosciences, 2019). The microbiota analyses were modified from a previously reported bioinformatics pipeline that incorporates the DADA2 protocol (95). The quality-trimmed reads were used to infer ribosomal sequence variants and their relative abundance in each sample after removing chimera. Taxonomy was assigned to each ASV generated by DADA2 using both the SILVA (release 132) database and Genome Taxonomy Database (GTDB) (96) and the RDP naïve Bayesian classifier as implemented in the *dada2* R package (97, 98). In a few cases when conflicted taxonomic assignments appeared, NCBI Refseq 16S rRNA combined with RDP database (99, 100) and Human Intestinal 16S rRNA database (HITdb v1) (101) were used to resolve the conflict. Pacific Biosciences long-reads sequencing complements short-reads sequencing for its high accuracy and extended length. To boost taxonomy assignment for short sequencing, we performed BLASTN search of the short-read ASVs to the long-read ASVs, and assigned the taxonomic name to the short reads if there is 100% percent identity and unanimous assignment if there are multiple hits to long-reads sequences.

A heatmap was constructed from the 50 most abundant intestinal bacterial taxa relative abundance in samples collected from 113 preterm infants enrolled in the study. The ASVs were normalized using total sum to calculate their relative abundances. Ward linkage clustering was used to cluster samples based on their Jensen-Shannon distance calculated in vegan package in R (102). The number of clusters was validated using gap statistics implemented in the *cluster* package in R (103) by calculating the goodness of clustering measure. Package raxml (v8.0.0) (104) was used to construct the phylogeny, Phyloseq R package (v1.38.0) (105) was used to display the phylogeny and the barplot. Volatility plot to demonstrate the fluctuation of microbial community diversity (characterized as Shannon diversity index) over MOM feeding volume in high or low IP groups. Plot was generated in QIIME (2019.10 vers) (106) (option -longitudinal plot-feature-volatility).

### Statistical analysis of intestinal microbial community

Hilbert-Schmidt Independence Criterion (HSIC) R package ‘dHSIC’ (107) was used to examine the independence between any variables with IP. Longitudinal modeling was performed using zero-inflated negative binomial random effects (ZINBRE) models. These models account for the possibility of existence of more than expected zeros (from negative binomial distribution) as well for correlations between samples from the same subject. Though IP was categorized to high and low groups, it is inherently continuous and hence we modeled IP as continuous value in our analyses. Subject was included as a random factor. Read counts data of phylotypes detected in at least 15% samples were modeled using ZINBRE models. The same principle was applied to MOM and PMA. The model was fitted using JAGS R package (108), and 10,000 iterations with the same number of burn in iterations was used. The convergence of the model was assessed using Gelman and Rubin’s potential scale reduction factor (109) and visual inspection of each coefficient’s Markov chains. The mean of the posterior distributions of estimated coefficients and their corresponding 95% credible intervals were calculated using model’s Markov chains. The credible intervals without overlapping are considered significant. P values were computed assuming normality of the posterior distributions of the corresponding coefficients. An adaptive spline logistic regression model implemented in spmrf R package (110) was used independently to confirm the association between B. breve to IP and MOM. This model is a locally adaptive nonparametric fitting method that operates within a Bayesian framework, which uses shrinkage prior Markov random fields to induce sparsity and provides a combination of local adaptation and global control (110). Bayesian goodness-of-fit p-value implemented in R package rstan (111) was used to access the significance of the association. R code implementation of the model is deposited in https://github.com/igsbma/IP_microbiome/tree/main/statistical_analyses. Discriminatory machine learning schemes computation were implemented in weka (112, 113), including J48 decision tree, REPTree, decision stump, and logistic model trees. The functional enrichment test was performed for each functional group (based on COG and PFAM annotation) and each of homologous gene cluster (HGC) generated in genome comparison analyses. The frequency tables of each function or HGC in each category (*i.e.*, MAGs of this study versus genBank genomes) were generated, which was used to fit a generalized linear model with the logit linkage function to compute an enrichment score and p-value for each unit (114). False detection rate correction to p-values was used to account for multiple tests using R package ‘qvalue’ (115).

### Intestinal microbiome analyses

Metagenomic sequence data were pre-processed using the following steps: 1) human sequence reads and rRNA LSU/SSU reads were removed using BMTagger v3.101 (116) using a standard human genome reference (GRCh37.p5) (117); 2) rRNA sequence reads were removed *in silico* by aligning all reads using Bowtie v1 (118) to the SILVA PARC ribosomal-subunit sequence database (94). Sequence read pairs were removed even if only one of the reads matched to the human genome reference or to rRNA; 3) the Illumina adapter was trimmed using Trimmomatic (119); 4) sequence reads with average quality greater than Q15 over a sliding window of 4 bp were trimmed before the window, assessed for length and removed if less than 75% of the original length; and 5) no ambiguous base pairs were allowed. The taxonomic composition of the microbiomes was established using MetaPhlAn version 2 (120). Metagenome assembled genomes (MAGs) pipeline includes *de bruijin* genome assembly using SPAdes v.3.10.1 (121), the bins were defined through distance clustering based on coverage and tetranucleotide signature using MetaBat v2 (122), and were refined using GTDB-Tk (89). Genomes were annotated using PROKKA v1.13 (90), annotated through evidences from the nomenclature of the consortium for function glycomics, eggNOG (v4.5)(123), KEGG 2013-03-18 release (124)), Pfam (v30.0)(125), CAZy (2014 release) (126, 127).

Similarity searches were performed to compare with previously annotated enzymes or transporter proteins based on the accession number (36–38), using BLASTP and confirmed with the COG, Pfam and CAZy annotation evidence to ensure the integrity of the results. The 8 essential extracellular enzymes that are known to be required in extracellular cleavage of HMOs before importing selected products of degradation are investigated (36–38), include: 1,2-α-l-Fucosidase (AfcA), 1,3/4-α-l-Fucosidase (AfcB), 2,3/6-α-Sialidase (SiaBb2), Lacto-N-biosidase (LnbB, LnbX), Chaperon for LnbX (LnbY), β-1,4-Galactosidase (BbgIII), β-N-Acetylglucosaminidase (BbhI). Five essential bacterial ABC transporters and homologs involved in the import of oligosaccharides were examined, which was known to show an exquisite specificity conferred by substrate-binding protein (SBPs) for different HMO molecules (39), including GNB/LNB (galacto-N-biose/lacto-N-biose I) transporter SBP (GltA), FL transporter SBPs (FL1-BP, FL2-BP), and LN*n*T transporter SBP (NahS). In addition to similarity search on *Bifidobacterium* genomes and MAGs, we also confirmed the results by searching the metagenomic community gene content, so to verify the target genes are not from species other than *Bifidobacterium*.

Metapangenomes were prepared using the MAGs constructed in this study and publicly available genomes under the species name *B. breve* (taxID: 1685) and *B. longum* (taxID: 216816), listed in **Table S6**. The metapangenome was constructed using anvi’o vers 6.2 (128) following pangenome workflow (114). Homologous gene clusters (HGCs) were identified in this set of genomes based on all-versus-all sequence similarity. Briefly, this workflow uses BLASTP to compute ANI identity between all pairs of genes, uses the Markov Cluster Algorithm (MCL) (129) to generate homologous gene clusters and aligns amino acid sequences using MUSCLE (130) for each gene cluster. Each gene was assigned to core or accessory according the hierarchical clustering of the gene clusters. Sourmash vers 3.3 (131) was used to compute ANI across genomes. To count as being present in the sample, it had to be at least 50 reads mapping on at least one *Bifidobacterium* species genomes, and the total abundance had to be at least 0.1% after normalizing over the total number of reads. For long-read data sequenced on Pacific Biosciences Sequel II platform, QC and assembly was performed using Canu-1.8 (86). The assemblies were assigned species name through BLAST to refseq dataset and confirmed with GTDB-Tk v1.1.0 (89). Genome alignment of the assemblies assigned to *B. breve* was aligned to reference *B. breve* genome JCM1192 using MAUVE aligner (132, 133).

### Data and Code Availability

All metagenomicm, metataxonomic and genomic data were deposited under BioProject PRJNA774819 (https://www.ncbi.nlm.nih.gov/bioproject/PRJNA774819) for open assessment. Illumina 16S rRNA V3V4 gene amplicon and Pacific Biosciences full-length 16S rRNA gene data were deposited in Sequence Read Archive with experiment ID from SRX12805867 to SRX12806634. Data deposition includes samples of positive and negative controls in each plate. Metagenomic data using Pacific Biosciences were deposited in SRR16598000 and SRR16598001. Metagenomic data using Illumina platform were deposited in the same BioProject with experiment ID from SRX12798907 to SRX12798933. The assembled genomes of *B breve* were deposited under the accession ID JAJGBR000000000 and JAJGBS000000000. The R code processing these sequences and SAS code used in this statistical analysis are deposited in https://github.com/igsbma/IP_microbiome/tree/main/statistical_analyses. Detailed information of sequences and annotation of pangenome can be retrieved at https://github.com/igsbma/IP_microbiome/tree/main/pangenome.

## SUPPLEMENTAL MATERIALS

FIGURE S1, PDF file

FIGURE S2, PDF file

FIGURE S3, PDF file

FIGURE S4, PDF file

FIGURE S5, PDF file

FIGURE S6, PDF file

TABLE S1, Excel file

TABLE S2, Word file

TABLE S3, Excel file

TABLE S4, Excel file

## CONTRIBUTIONS

B.M., J.R., S.S., and R.V. designed the research; S.S., E.J., E.M., B.M., G.N., H.Y., L.S.R. and R.V. conducted the clinical study; B.M., H.Y., L.F., and M.H. conducted the short-read sequencing; B.M., E.M., M.H., L.S., L.J.T. conducted the long-read sequencing; B.M., M.F, and P.G. performed the statistical analyses; B.M. and G.N. performed the epidemiological analyses; B.M., S.S., J.M.L-D., G.N., M.F., M.F.P., J.R., and R.V. wrote the paper.

## ACKNOWLEDGEMENTS

This study was supported in part by The Gerber Foundation 2018 award (Project ID 6361), the National Institute of Diabetes and Digestive and Kidney Diseases (NIDDK) of the National Institute of health under award number R21DK123674, and the Institute for Clinical and Translational Research (ICTR) at University of Maryland Accelerated Translational Incubator Pilot (ATIP) award. The authors thank Dr. Jonathan Meddings and Mr. Kim Le at the University of Calgary, Calgary, Alberta, Canada for the HPLC analysis of serum and urine samples. Ms. Ivette Santana-Cruz (GRC at IGS) for constructive discussion on long-read data processing. Ms. Shilpa Narina and Sarah Arbaugh for great assistance in clinical specimen collection.

## Supplementary Figures and Table legends

**FIG S1** Intestinal permeability (IP) and cohort clinical information. **A**) Notched boxplot of IP for early preterm subjects (GA < 33 weeks gestation). Subjects were categorized by IP-measuring day between study day 7-10 and by IP category. The top and bottom of the box are the lower and upper quartiles, and the band near the middle of the box represents the median. The width of the notch can be used to roughly compare two distributions. For example, two distributions without overlapping notch regions can be roughly considered as being significantly different from each other (1). IP was measured by non-metabolized sugar probes lactulose and rhamnose. High IP was defined by a La/Rh ratio >0.05, as validated and applied previously (2). **B**) Correlation matrices visualization of the subjects’ physiological age. R package Correlograms (corrgram) were used to visualize the correlation matrices. Pearson correlation method used to calculate correlation. **Abbr**: PMA at dosing: postmenstrual age calculated at the dosing day when IP was measured; PMA at enrollment: postmenstrual age at enrollment day taken place within 1-4 days post-birth; GA: gestational age; BW: birthweight; body weight at dosing: subject weight measured at the dosing day when IP was measured.

**FIG S2** Phylogenetic tree of all ASVs of full-length 16S rRNA gene sequenced on Pacific Biosciences Sequel II platform. Package raxml (v8.0.0) (3) was used to construct the phylogeny, Phyloseq R package (4) was used to display the phylogeny.

**FIG S3** Heatmap of the 50 most abundant intestinal bacterial taxa relative abundance in samples collected from 113 preterm infants enrolled in the study. The fecal microbiota was characterized by high-throughput sequencing of the V3-V4 variable regions of 16S rRNA genes. Ward linkage clustering was used to cluster samples based on their Jensen-Shannon distance calculated in vegan package in R (5). The number of clusters was validated using gap statistics implemented in the *cluster* package in R (6) by calculating the goodness of clustering measure.

**FIG S4** Information on bifidobacterial abundance and intestinal permeability (IP). **A**) Relative abundance of bifidobacterial bacterial groups stratified by feeding types. Phyloseq R package (v1.38.0) (4) was used to generate the barplot. **B**) The relative abundance of *B. breve* between high-IP and low-IP groups. Dependence between **C**) IP or **D**) MOM feeding dose and the log relative abundance of *B. breve*. An adaptive spline logistic regression model implemented in spmrf R package (7) was applied to the phylotypes present in at least 15% of all samples. Bayesian goodness-of-fit p-value implemented in R package rstan (8) was used to access the significance of the association between phylotypes and investigated factors.

**FIG S5** Metapangenome of *Bifidobacterium breve*. The 26 *in-house B. breve* MAGs was supplemented with 107 published genomes (https://doi.org/10.6084/m9.figshare.19709917.v2 **A**) and our 4 *B. longum* MAGs was supplemented with 310 published genomes (https://doi.org/10.6084/m9.figshare.19709917.v2 **B**) for pangenome construction following pangenome workflow (9). *B. breve* pangenome was displayed using anvi’o *vers* 6.2 (10). BLASTP was used to compute ANI identity between all pairs of genes. Markov Cluster Algorithm (MCL) (11) was used to generate homologous gene clusters (HGCs). Amino acid sequences of each HGC were aligned using MUSCLE (12). HCG was assigned to core, accessory or dispensable according the hierarchical clustering of the gene clusters. Detail of each HGC was in https://doi.org/10.6084/m9.figshare.19709917.v2 **C**. Sourmash vers 3.3 (13) was used to compute Average nucleotide identity (ANI) across genomes. The source indicates the isolated origin of the genome, and genomes of the same subject are indicated in the same cohort.

**FIG S6** The complete *B. breve* genome reconstructed in this study. Metagenomic sequencing of the two selected fecal samples was performed using the Pacific Bioscience Sequel II platform, followed by assembly using Canu v1.8 (14) and deconvolution using BLASTN of the assembly. This complete genome was 2.34M in size (https://doi.org/10.6084/m9.figshare.19709923.v1, https://doi.org/10.6084/m9.figshare.19723255.v1 **C**), similar to median *B. breve* genome size of 2.33M reported on NCBI. **A**) KEGG 2013-03-18 release (15) to characterize functional categories of *B. breve* XM1439. **B**) Circular genome display of *B. breve* XM1439, generated by BLAST Ring Image Generator (BRIG) (2011 June vers) (16). **C**) Genome alignment of *B. breve* genome 1439, 1437 using MAUVE (17) using *B. breve* DSM20213 as the reference genome.

**TABLE S1.** Clinical metadata of the 113 early preterm infant subjects used in this study.

**TABLE S2.** Dependence of demographic, obstetric, and neonatal characteristics with intestinal permeability (IP) using Hilbert-Schmidt Independence Criterion (HSIC) implemented in R package dHSIC.

**TABLE S3.** Taxonomic groups significantly associated with PMA, IP and MOM feeding volume. Zero-inflated negative binomial random effects (ZINBRE) models were used to compute significance level of association, which accounts for many zeros as well for correlations between samples from the same subject. All phylotypes detected in at least 15% samples were modeling using ZINBRE models. PMA, IP and MOM feeding volume were modeled as continuous value. **A**) Taxonomic groups associated with PMA, which was calculated as day of life after birth plus gestational age; **B**) Taxonomic groups associated with IP, measured at 7-10 days after birth; **C**) Taxonomic groups significantly associated with MOM feeding volume. **Abbr**: **MOM**: mother’s own breastmilk feeding. **PMA**: postmenstrual age. **IP**: intestinal permeability.

**TABLE S4.** *Bifidobacterium* homologous gene clusters (HGCs) characterized to involved in human milk oligosaccharides assimilation. Genomes were annotated through annotative evidences from the nomenclature of the consortium for function glycomics, eggNOG (v4.5)(18), KEGG (FTP Release 2013-03-18)(15)), Pfam (v30.0)(19), CAZy (2014 release) (20, 21). Similarity searches were performed to previously annotated enzymes or transporter proteins based on the accession number listed in previous studies (22–24), using BLASTP similarity search and confirmed with the COG, Pfam and CAZy annotation evidence to ensure the integrity of the results. **A**) HGCs involved in extracellular enzymes and their homologs involved in extracellular cleavage of HMOs; **B**) HGCs characterized as family 1 solute binding proteins (F1SBP); **C**) HGCs involved in enzymes for catabolizing HMOs substrates intracellularly; **D**) HGCs characterized as FHMO (Fucosylated Human Milk Oligosaccharide utilization cluster); **E**) HGCs involved in sialylated HMO substrates catabolism; **F**) HGCs involved in sulfatase catabolism activity.

